# Conformal Prediction of Molecule-induced Cancer Cell Growth Inhibition Challenged by Strong Distribution Shifts

**DOI:** 10.1101/2024.03.15.585269

**Authors:** Saiveth Hernandez-Hernandez, Qiarong Guo, Pedro J. Ballester

## Abstract

The drug discovery process often employs phenotypic and target-based virtual screening to identify potential drug candidates. Despite the longstanding dominance of target-based approaches, phenotypic virtual screening is undergoing a resurgence due to its potential being now better understood. In the context of cancer cell lines, a well-established experimental system for phenotypic screens, molecules are tested to identify their whole-cell activity, as summarized by their half-maximal inhibitory concentrations. Machine learning has emerged as a potent tool for computationally guiding such screens, yet important research gaps persist, including generalization and uncertainty quantification. To address this, we leverage a clustering-based validation approach, called Leave Dissimilar Molecules Out (LDMO). This strategy enables a more rigorous assessment of model generalization to structurally novel compounds. This study focuses on applying Conformal Prediction (CP), a model-agnostic framework, to predict the activities of novel molecules on specific cancer cell lines. A total of 4320 independent models were evaluated across 60 cell lines, 5 CP variants, 2 set features, and training-test splits, providing strong and consistent results. From this comprehensive evaluation, we concluded that, regardless of the cell line or model, novel molecules with smaller CP-calculated confidence intervals tend to have smaller predicted errors once measured activities are revealed. It was also possible to anticipate the activities of dissimilar test molecules across 50 or more cell lines. These outcomes demonstrate the robust efficacy that LDMO-based models can achieve in realistic and challenging scenarios, thereby providing valuable insights for enhancing decision-making processes in drug discovery.

## 1 Introduction

The drug discovery process typically starts with identifying a small organic molecule with the potential to become a drug candidate (1). There are two main strategies: phenotypic virtual screening (VS) and target-based VS. In target-based VS, a defined molecular target has been validated for the considered disease and is leveraged to find molecules with activity for that therapeutic target (2). For instance, crystal structures of the target can be exploited to identify such active molecules (3,4). In contrast, phenotypic drug discovery (PDD) does not rely on a specific molecular target or its mechanism of action in treating the disease (2,5). For instance, PDD leverages molecule-induced phenotypic changes in a cell (6). Despite target-based drug discovery being the predominant approach for the past 30 years, PDD has been shown to be more effective than it has been given credit for (7–9) and, hence, has been undergoing a resurgence (5,6). Cancer cell lines are a well-established experimental system for PDD screens (10). Molecules are tested on a cell line to identify their sensitivity or resistance, as determined by their half-maximal inhibitory concentrations (11). These concentrations are denoted as IC_50_s or GI_50_s or their logarithmic versions, pIC_50_s or pGI_50_s.

Machine Learning (ML) has become a powerful approach to building models for phenotypic VS (12,13). Investigating the most suitable learning algorithms for this and related purposes is an active research area (14–16). In the context of cancer cell lines, research has been intense on the precision oncology side (17–19), where the activities of cell lines when treated with a given drug molecule are predicted. However, studies for phenotypic VS (20,21), where the activities of molecules on a given cell line are predicted instead, are still scarce.

Improving generalization remains a critical step toward building robust predictive ML-based models (22). A limited generalization of such models to unseen molecules often arises from biases in the way training and test sets are constructed, such as through random or scaffold-based splitting strategies, which may not adequately represent the diversity or complexity of real-world screening scenarios (23). A promising direction to address this challenge is by leveraging clustering for model validation. The intuition is that molecules from different clusters should have much lower similarity than molecules from the same cluster. In prior work (24), we systematically evaluated multiple clustering algorithms to identify an optimal clustering configuration tailored to the National Cancer Institute (NCI-60) dataset (10). The resulting clusters are then used in a Leave Dissimilar Molecules Out (LDMO) cross-validation approach, where by holding out entire clusters for testing, we can evaluate model performance on genuinely novel chemical series, thereby providing a more realistic and rigorous benchmark for generalization.

Equally important is the ability to quantify predictive uncertainty, not only to assess the trustworthiness of individual predictions, but also to prioritize candidates for experimental validation in a cost-effective manner and reduce the risk of pursuing false positives (21). However, reliability is usually assigned to the whole model (e.g., as the RMSE between predicted and measured activities of test set molecules) rather than reliability at the instance level (e.g., as a predicted activity interval where the measured activity of the corresponding test set molecule is most likely to be). In this context, conformal prediction (25) is a model-agnostic framework for generating prediction sets with user-defined error rates. Therefore, the application of CP to several drug design problems has been investigated (26–29). In particular, we published a proof-of-concept study showing that CP improves phenotypic VS on cancer cell lines on random training-test dataset splits (21). Although CP is a promising tool, it still relies on the validation strategy. Therefore, cross-conformal prediction or CCP (30) has been proposed as an extension of the standard CP framework, which mitigates the sensitivity of CP to a single calibration set by averaging nonconformity scores across multiple folds.

In this work, we address both generalization and uncertainty quantification by introducing LDMO-CCP, a clustering-based validation framework for conformal prediction. Here, we study how well these CP models perform in a much more realistic and challenging scenario where the test molecules are not only unseen but also dissimilar to those in the training set. At the initial stage, a total of 2100 models (60 cell lines x 5 CP models x 7 clusters) are built and evaluated. The two best-performing CP models advance to the next stage, resulting in the evaluation of 120 models (60 cell lines x 2 CP models). Since evaluations occur for each cell line, the results remain robust, derived from the analysis of 2220 independent cases. We investigate how different are the chemical structures of the most potent cell-active molecules that were correctly predicted. We also assess these models’ potential to anticipate the activities of a dissimilar test molecule across cell lines.

We demonstrate that the LDMO-CCP approach enables more reliable evaluation of model performance and better highlights the uncertainty associated with predictions on novel chemical space, offering practical improvements for real-world compound screening efforts. Figure 1 summarizes the proposed pipeline for narrowing the NCI-60 dataset to a manageable subset of candidate molecules for further testing, such as *in vitro* assays.

**Figure 1.**
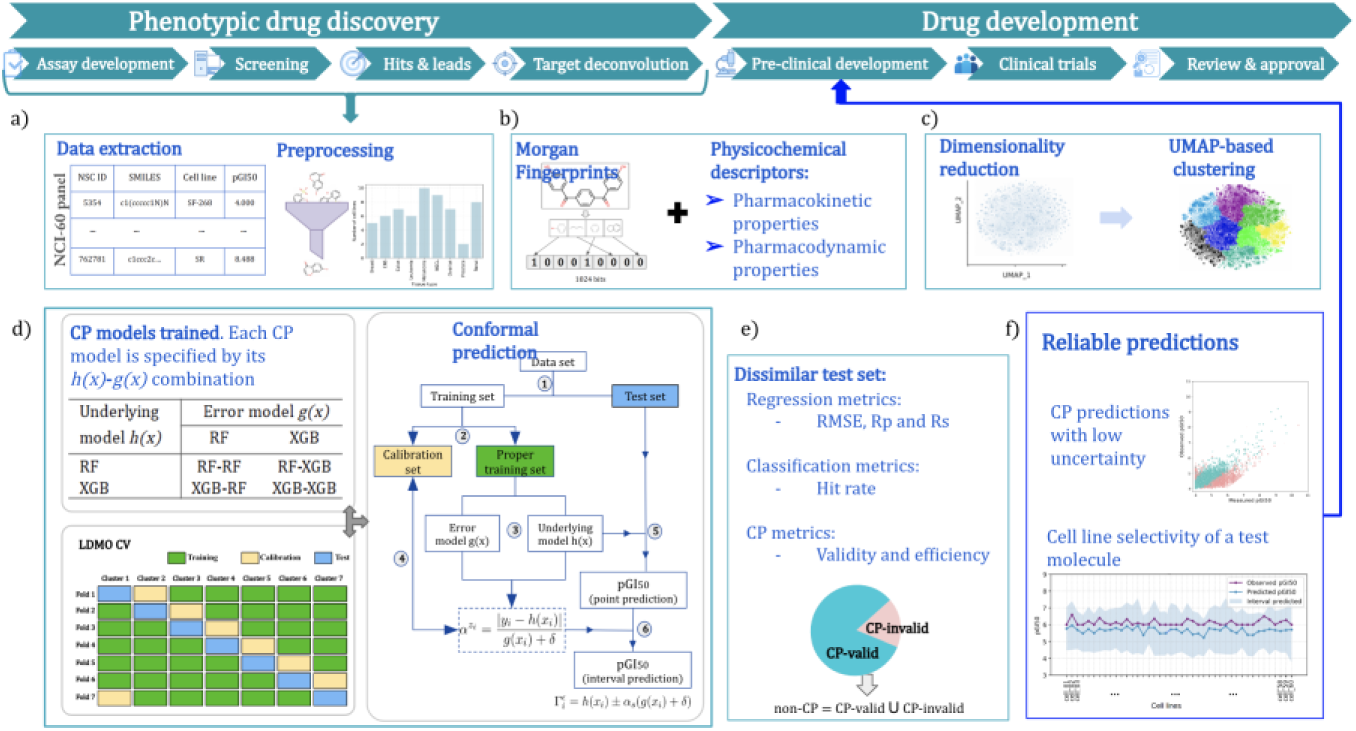
Proposed pipeline for narrowing down the NCI-60 dataset to a manageable subset of candidate molecules with a high likelihood of biological activity by using the LDMO-CPP approach. a) Molecules from the NCI-60 panel are extracted and preprocessed. b) Generating the features for each molecule using MFPs and PCDs to encode their molecular characteristics. c) Molecular clustering of the unique molecules in the NCI-60 dataset, regardless the cell line. For each of the 60 cell lines: d) For each model, multiple LDMO-CV validation runs are performed to evaluate how well the considered model can predict the cell activity of molecules with different chemistry. e) The standard ML metrics and the CP metrics are used to evaluate model performance. f) Reliable predictions can be then tested in pre-clinical trials, such as *in vitro* assays.

## 2 Material and Methods

### 2.1 Datasets

The availability of preclinical data enables the development and validation of ML-based models in drug design. Such is the case of the National Cancer Institute (NCI-60) human tumor cell lines screen, which since 1990 has been used by the cancer research community to find compounds with potential anticancer activity. The NCI-60 (10) panel utilizes 60 different human tumor cell lines to identify and characterize novel compounds with growth inhibition or killing of tumor cell lines. These 60 different human tumor cell lines comprise 9 broad cancer types: leukemia (LE), melanoma (MEL), lung (NSCL), colon (CO), brain (CNS), ovary (OV), breast (BR), prostate (PR), and kidney cancers (RE). All compounds screened are initially tested in a one-dose assay on the full NCI-60 panel. Compounds showing significant growth inhibition at this assay are then evaluated against the NCI-60 panel in a 5-dose assay (10).

The results of these experiments are available for download from the DTP Bulk Data for Download website, in comma-delimited file format (Figure 2). Each compound screened by the NCI-60 is identified with a unique registration number called the National Service Center (NSC) ID. In this study, we modeled the pGI_50_ of molecule-cell line pairs, defined as the negative logarithm of the half-maximal inhibitory concentration of the molecule on the cell line.

**Figure 2.**
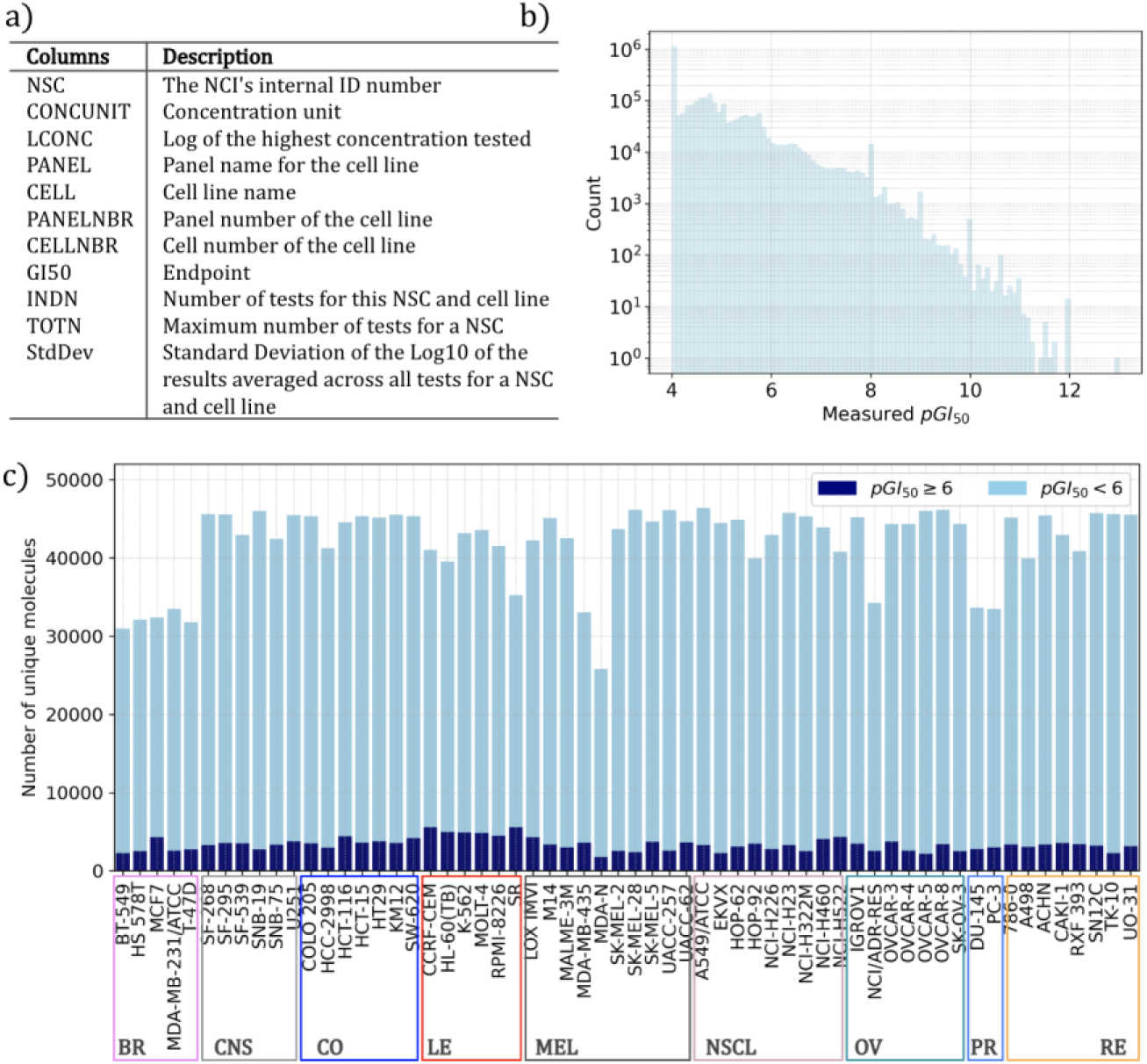
NCI-60 Growth Inhibition Bulk Data description. a) The NCI-60 Growth Inhibition Data is available for download in comma-delimited file format and includes 11 fields that describe the data. b) Distribution of the *pGI*_50_ measurements in the molecules across the 60 cell lines for a total of 2,707,434 data points, where potent molecules (*pGI*_50_ ≥ 6) are rather scarce. c) Distribution of the number of unique molecules tested per cell line (60 distinct cell lines representing nine different types of cancer). A color code is assigned to show the number of potent molecules (*pGI*_50_ ≥ 6) per cell line. The x-axis also shows the cell lines for each of the nine cancer types.

Data quality plays a key role in developing AI/ML models. Therefore, data cleansing is necessary to ensure the use of high-quality data in generating these models. To achieve this, we follow the preprocessing protocol described by some of us (21) and comprising the removal of low-activity (pGI_50_ < 4) molecules as they do not have therapeutic potential and the calculation of the mean when more than one pGI50 measurements were available for the same NSC-cell line pair. Next, each molecule is represented by its 1024-bit MFP with a radius of 2, corresponding to the presence or absence of a particular substructure in the molecule. The 1024-bit MFP size has been successfully used in retrospective studies, e.g., in Siramshetty et al. 2020 (31). While the choice of fingerprint and metric typically has little effect on downstream predictions. For example, in target prediction based on molecular similarity (32), this choice led to practically the same performance as other fingerprints and metrics, suggesting that it should be a near-optimal choice here. This process obtained 2,707,434 data points (50,555 small molecules screened against 60 cancer cell lines), representing a matrix completeness of 89.25%.

For these 50,555 small molecules, we also computed seven PCDs to encompass both the pharmacokinetic and pharmacodynamic aspects of drug action. These PCDs include molecular weight, logP, number of hydrogen bond donors (HBD), number of hydrogen bond acceptors (HBA), total polar surface area, number of aliphatic rings, and number of aromatic rings. These descriptors provide insights into the molecular characteristics influencing the absorption, distribution, metabolism, and excretion or ADME processes (pharmacokinetics), as well as the interaction of the drug with its molecular targets (pharmacodynamics). These features were calculated using the RDKit software suite (33).

### 2.2 Learning algorithms

Selecting an appropriate learning algorithm is important in achieving generalization in ML models. While deep learning (DL) approaches have demonstrated strong performance in similar applications (34,35), recent literature has shown that tree-based models frequently outperform DL approaches on tabular data (36). Therefore, considering the NCI-60 dataset used in this study and the strong performance of tree-based models in our previous work using random validation strategies (21), here we use the Random Forest (RF) and eXtreme Gradient Boosting (XGB) algorithms. In addition, we include Linear Regression (LR) as a baseline to evaluate the benefits of moving from linear to non-linear modeling approaches.

These algorithms have been extensively documented in prior literature; however, we provide a summary of their key characteristics for accessibility. The LR algorithm is used here as a performance baseline (37). LR fits a linear model to minimize the difference between observed and predicted values across training instances, using the features as independent variables. RF algorithm generates an ensemble of diverse tree predictors that collectively constitute a nonlinear model (38). In regression, RF predicts an instance’s label as the average of the individual tree predictions. The XGB algorithm weights the prediction of each generated tree by how well it individually matches the data (39). XGB was designed to optimize training speed and generalization performance. We use the scikit-learn implementations (40) for these algorithms.

### 2.3 Performance metrics

The evaluation of the CP models involves the use of regression and classification ML metrics, which include the Root Mean Squared Error (RMSE), Pearson correlation (Rp), Spearman correlation (Rs), and hit rate (23). The RMSE measures the average magnitude of errors between predicted and actual values, where lower RMSE values indicate better model accuracy, with 0 being a perfect fit. The Rp quantifies the linear relationship between predicted and actual values. It ranges from –1 to 1, indicating perfect negative and positive correlations. The Rs assesses the monotonic relationship between predicted and actual values. It ranges from –1 to 1, with values close to 1 or –1 implying a strong monotonic correlation, while 0 indicates no monotonic relationship.

To calculate the hit rate (HR), we establish a threshold of 6 pGI_50_ units (28) to determine whether predictions are positive (active) or negative (inactive). A prediction is positive if it is higher than or equal to 6 (y_i_ ≥ 6); otherwise, it is negative (y_i_ < 6). The HR measures the proportion of true positives in compounds identified as positive. This allows us to prioritize compounds, aligning with the objective of virtual screening to identify potential leads.

In addition, we employ efficiency and validity metrics from the CP framework (25,28) to complement our evaluation. Validity measures the reliability of the predictions, indicating that at a confidence level of 80% (0.8), the predictor will include the actual value within its prediction intervals at least 80% of the time, assuming CP is well-calibrated. Efficiency, on the other hand, measures the specificity of the predictions and is related to the interval size (uncertainty). A higher confidence level reduces efficiency, resulting in larger predicted intervals, i.e., predictions with higher uncertainty.

### 2.4 Hyperparameter tuning

Hyperparameter tuning is a pivotal strategy to augment the efficacy of ML algorithms, especially when applied to complex problem domains such as the prediction of inhibitory activity of molecules on a cancer cell line. In our previous work (21), we performed hyperparameter tuning to find the best combination of the most critical hyperparameters for the RF and XGB algorithms using a systematic grid search approach. More concretely, we did this for each of the 60 labeled molecule datasets, one per cell line, adhering to an 80/10/10 partitioning scheme for training, validation, and test sets. In our current work, we are utilizing the optimal hyperparameter values that we have identified in our previous work, as for each cell line, multiple models are trained as a part of the LDMO CV and outlier-based partitions.

## 3 Methodology

### 3.1 Conformal Prediction

Conformal prediction, or CP is a flexible framework that can be applied to various ML algorithms without requiring modifications to the underlying model (25). Unlike conventional predictive models that provide single-point predictions, CP creates prediction sets that include the measured value with a user-specified confidence level (26,28).

Given a training data set *(x*_1_*,y*_1_*), (x*_2_*,y*_2_*), …, (x*_n_*,y*_n_) and a new test instance *x*_n+1_, CP defines the nonconformity score *α* for each point based on how well the model’s predictions align with the observed values. For regression problems, the nonconformity score is often a function of the residuals (26). Thus, CP works by predicting that a new test instance will have a label similar to the labels of old instances (calibration set) based on a specified similarity measure (calibration function). The objective is to create subsets, *Γ(x*_n+1_), of Y that are sufficiently large for the true label to fall within the subset (Figure 3).

**Figure 3.**
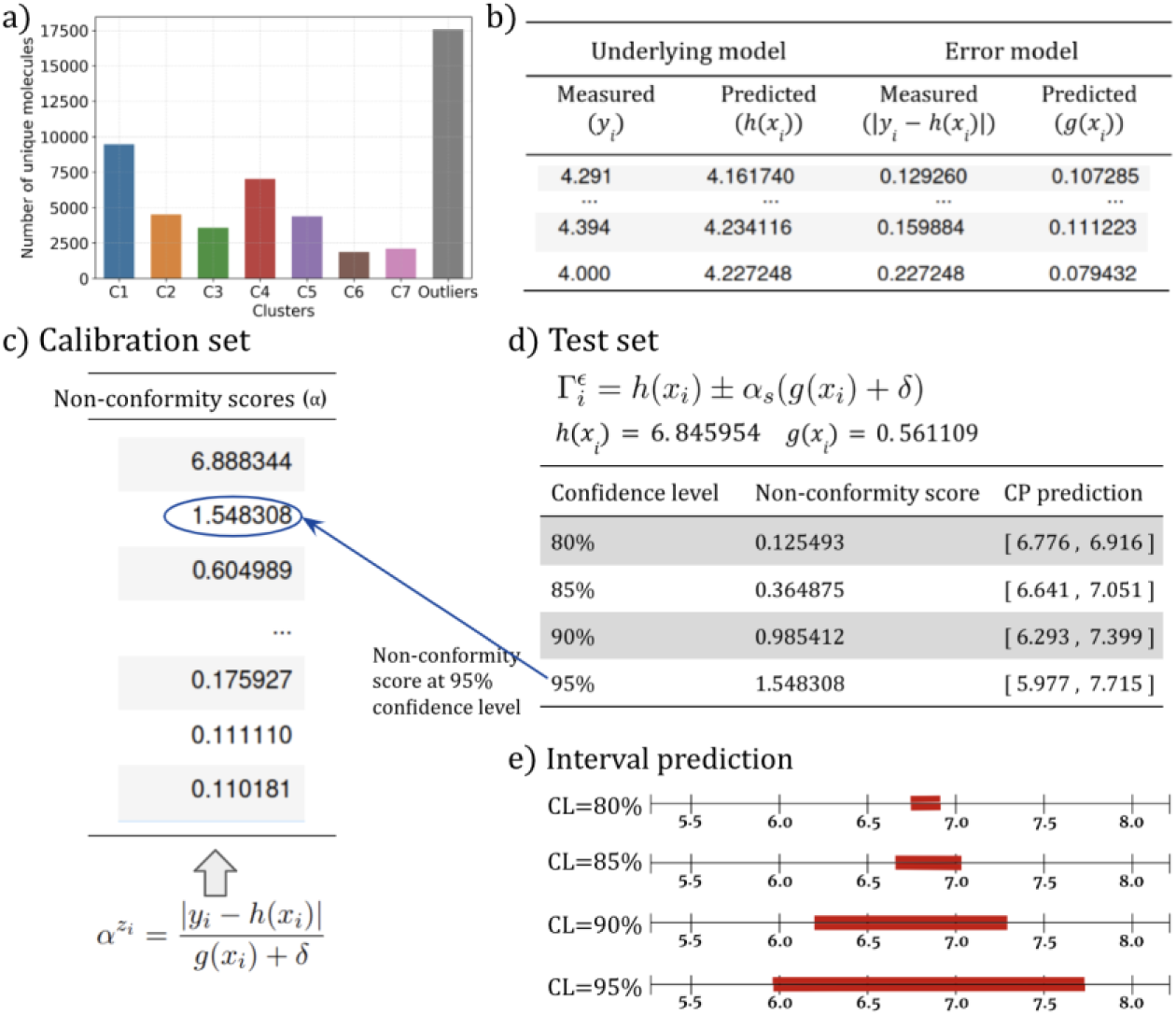
Conformal Prediction enhances an underlying model with reliability estimates at a given confidence level. a) Clusters used to build a LDMO-CCP model per cell line. At each fold, 5 clusters are used as a proper training set, 1 cluster as a calibration set, and the remaining 1 cluster is used as a test set. b) At each fold, two supervised learning models are trained on the proper training set: the underlying model, ℎ(*x*), and the error model, g(x). ℎ(*x*) is trained on y, here pGI_50_s, and *g*(*x*) is trained on the resulting error residuals (|*y* − ℎ(*x*)|) from the proper training set. c) The non-conformity scores are calculated for each sample, *Z*_*i*_ = (*x*_*i*_, *y*_*i*_), in the calibration set, using the underlying and error models trained on the proper training set samples. Each score represents how far was the predicted error to match the actual error for that calibration set sample. d) Given an unseen instance *x*_*i*_ for which a prediction is desired, the underlying model generates a point prediction, such as ℎ(*x*_*i*_) = 6.846 in this example. Additionally, the error prediction is computed for this instance, in this example, *g*(*x*_*i*_) = 0.561. The non-conformity score is selected based on a user-defined confidence level. For instance, at a 95% confidence level, the non-conformity score is 1.548 because 95% of the scores in the calibration set are lower than this value. The interval prediction is formed by combining the predictions from both the underlying model ℎ(*x*) and the error model *g*(*x*), along with the non-conformity score. Thus, by constructions, with a confidence level of 95% the true value will be within the predicted confidence intervals in at least 95% of the cases. In retrospect, if the interval contains the true value, these predictions are called CP-valid, otherwise, they are called CP-invalid. E) Graphical representation of the interval predicted (red) at each confidence level. Note that *uncertainty* = 2α_*s*_*g*(*x*_*i*_).

### 3.2 Leave Dissimilar Molecules Out approach

Standard CP-based methods may have limitations when the distribution of test data differs significantly from the training data, leading to decreased prediction accuracy and reliability (22,41). Consequently, advanced CP techniques like cross-conformal prediction (CCP) are being explored to address these challenges (30,42). In CCP, the dataset is split into k non-overlapping sets, similar to the k-fold cross-validation (CV) technique. Each set is used as a separate hold-out set, with models being trained on the remaining data. This results in the training of k CCP models, with each model using a different set as the calibration set (43,44).

In this study, we proposed the Leave Dissimilar Molecules Out (LDMO) CV approach, a modified variant of CCP, where the folds are obtained through a previous clustering process (24). The LDMO validation underlines that the test set molecules are not only unseen but also dissimilar to those in the training set. Unlike random partitioning or scaffold partitioning, which can introduce bias if structurally similar molecules appear in both training and test sets (22,23), molecular clustering forces the model to confront novel chemical entities in the test folds. Therefore, the proposed LDMO-CCP approach is advantageous in real-world circumstances where data distributions may vary over time or across different environments, as it fosters confidence in the prediction performance.

### 3.3 UMAP-based split for each of the 60 datasets

Molecular clustering involves grouping similar molecules based on their chemical properties or structural features. In a previous study (24), we identified a UMAP-based clustering method as the best way to cluster the molecules screened by the NCI-60 panel. The molecules are encoded using MFPs of size 1024 and radius 2. UMAP-based clustering refers to first applying the UMAP dimensionality reduction algorithm and subsequently performing agglomerative clustering within the resulting embedded space.

In (24), we first defined an outlier as the molecule whose Tanimoto similarity to its most similar molecule is lower than or equal to a given cutoff. Thus, an outlier molecule differs from any other molecule in the set at the predefined outlier cutoff. As a result, 32,971 non-outlier and 17,584 outlier molecules were retrieved at an outlier cutoff of 0.5. Note that the proposed definition for detecting outlier molecules serves a purpose beyond simply improving clustering results by cleaning the data. Specifically, we leverage this outlier set to establish an external dataset for evaluating our LDMO-CCP models.

Utilizing the UMAP-based clustering method, 7 clusters were derived for non-outlier molecules, with the outlier set forming its distinct cluster. The outlier detection and clustering steps were applied to the entire NCI-60 panel, regardless of cell lines. This approach saves time and resources while ensuring consistent results across the dataset. It also enhances scalability, allowing for more molecules or cell lines to be included in future analyses without significant computational burden. Moreover, clustering is employed in our retrospective study to capture the relationship between a training set and a real-world prospective virtual screening library. As a result, no clustering is required during prospective application, avoiding additional computational overhead.

The clustering output includes NSC ID, SMILES representations, and assigned clusters, facilitating mapping to individual cell lines. Each of the 50,555 molecules receives a cluster ID from 1 to 7 or is marked as Outlier. Combining NSC-ID with the cluster ID allows mapping clustering results per-cell line, resulting in eight distinct clusters for each cell line.

Finally, clusters 1 to 7 (non-outlier set) are used to build and evaluate a CP model using the LDMO cross-validation strategy. In this stage, at each fold, 5 clusters are used as the proper training set, 1 as the calibration set, and 1 as the test set. In the second phase, the non-outlier set is used to retrain the models (6 clusters as the proper training set and 1 as the calibration set), which are subsequently evaluated on the outlier set. This approach guarantees that the test set includes molecules that differ significantly from those used for model development.

## 4 Experiments and results

We designed two scenarios to evaluate CP models on dissimilar molecules that simulate real-world screening conditions. First, we trained LDMO-CCP models using seven-fold cluster-based splits, ensuring distinct training, calibration, and test sets. At this stage, 4,200 models (60 cell lines × 5 CP models × 2 feature sets × 7 folds) were evaluated. In the second phase, the top two LDMO-CCP models were retrained on non-outlier data (clusters 1 to 7) and tested on outlier molecules, resulting in 120 additional models (60 cell lines × 2 CP models).

All experiments were conducted on a DELL Precision Tower 7920 workstation equipped with dual Intel Xeon Gold 6136 CPUs, 128 GB of system memory, and running Ubuntu 22.04 LTS. No GPU acceleration was used. On average, the CP-based models can screen 10,000 molecules in roughly 60 seconds, making it suitable for high-throughput virtual screening tasks involving large compound libraries.

### 4.1 Strong distribution shifts challenge non-CP model predictions in all 60 cell lines

We initially conducted a performance comparison of the LR, RF, and XGB models (*h(x)*) to determine the most informative molecular features for predicting pGI_50_. We assessed two feature sets: Morgan Fingerprints descriptors (MFPs) of 1024 bits and a combined set of Morgan Fingerprints and 7 Physicochemical Descriptors (MFPs+PCDs). The objective was to understand how the inclusion of PCD contributes to model accuracy, as measured by their pGI_50_ prediction error (RMSE) and correlation (Rp and Rs). To ensure the accuracy of the results, we used the LDMO-CV approach (clusters 1 to 7) to train and evaluate each of the 60 non-outlier datasets (one per cell line). The results of the LDMO CV run are summarized with the median of the model evaluation scores. Since the underlying model *h(x)* is a regression model, i.e., without CP, confidence levels are not applicable at this stage. Therefore, such predictions are referred to as non-CP predictions.

Figure 4 compares how well the two sets of features, MFPs, and MFPs+PCDs, perform in terms of RMSE, Rp, and Rs across 60 cell lines. We performed a statistical test to assess the significance of the difference in performance between the models trained and evaluated with these two sets of features. The results indicate that including PCDs as input features (MFPs+PCDs) leads to a statistically significant improvement in the correlation between observed and predicted pGI_50_ values, as indicated by the Rs and Rp metrics. This suggests that including the PCDs enhances the model’s robustness, allowing it to adapt more effectively to data distribution shifts. However, regarding RMSE, results indicate no significant difference between the two feature sets, regardless of the model used. The comparable RMSE values suggest that the model’s predictive accuracy in terms of absolute errors remains consistent across both feature sets. As a result, the MFPs+PCDs feature set was chosen for subsequent analysis.

**Figure 4.**
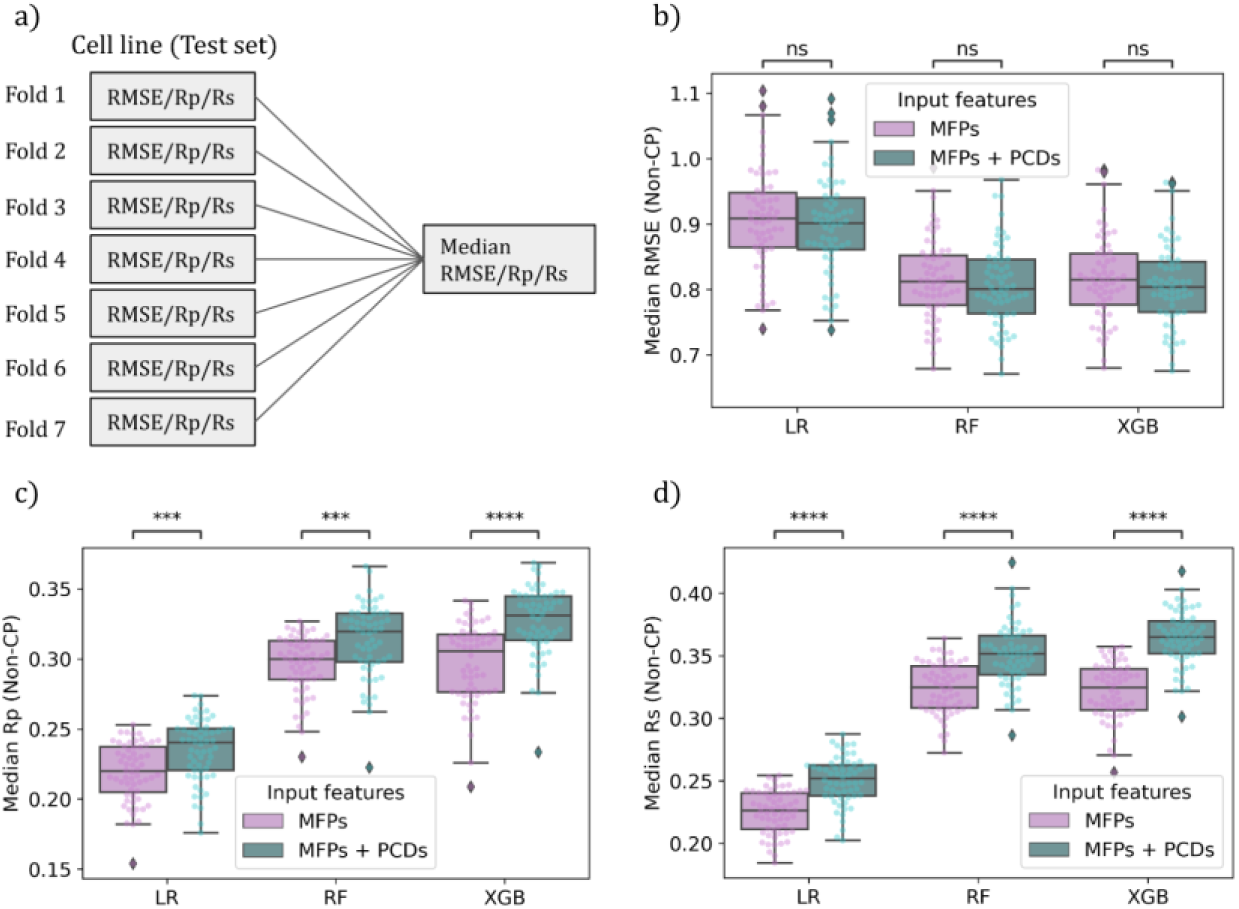
The three underlying models improve when the PCDs are included as part of the input features (MFPs+PCDs). Each molecule is encoded with an MFP of 1024 bits and 7 physicochemical descriptors. LR, RF, and XGB were used to evaluate the performance of each feature set (MFPs and MFPs+PCDs, identified with a color code), with each algorithm trained and evaluated on non-outliers molecules using an LDMO-CV approach (each fold being a different cluster of molecules). a) For each cell line, the evaluation score of the model is the median of the evaluation score of each fold, i.e. a 7-fold-median score. From b) to c), each boxplot represents, respectively, the distribution of the median RMSE, Rp, and Rs in predicting pGI_50_ values across the 60 cell lines. The results of the statistical test performed for the underlying models have been placed at the top of the boxplots. p-value annotation legend: ns: 5.00e-02 < p <= 1.00e+00; ***: 1.00e-04 < p <= 1.00e-03; ****: p <= 1.00e-04

In addition to this, results from Figure 4 demonstrate that non-linear models exhibited superior performance compared to the linear regression model. Based on this finding, the subsequent analyses were carried out exclusively with non-linear models.

### 4.2 CP models are substantially more accurate than non-CP models in this scenario

We evaluated four confidence levels: 80%, 85%, 90%, and 95%, corresponding to expected errors of 0.2, 0.15, 0.1, and 0.05, respectively. For regression problems, the expected errors indicate the quartile in which the nonconformity score will be selected (30,44). Figure 5 illustrates the uncertainty distribution for each confidence level, using dotted lines to depict the requested error value for each level. To calculate the uncertainty, we multiplied the predicted error for a specific molecule by the selected nonconformity score at a particular confidence level, expressed as uncertainty = 2αsg(x_i_).

**Figure 5.**
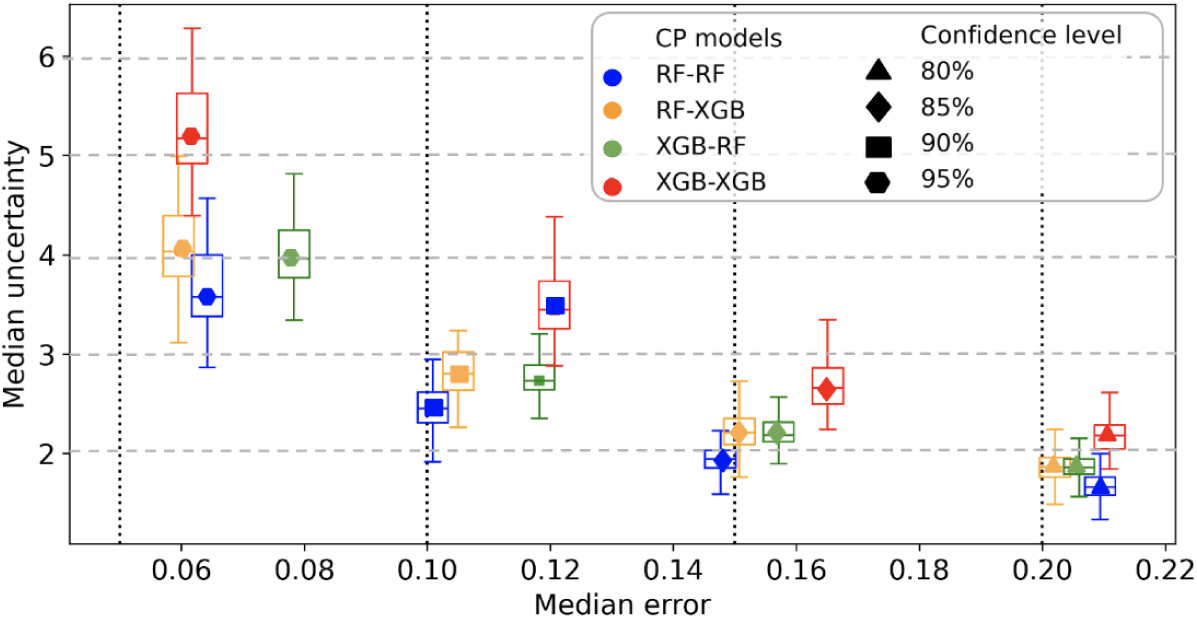
Trade-off between prediction uncertainty and measured error of CP models. For each CP model, median error and uncertainty are computed using MFPs+PCDs as input descriptors at different confidence levels. The cell lines’ performance is summarized as 7-fold-median uncertainty (y-axis). Thus, boxplots contain 60 points, representing the median uncertainty per cell line. The median of 7-fold-median error values is calculated across 60 cell lines to summarize the error performance. A single point is obtained that represents the median error of the entire group, and each boxplot is positioned along the x-axis based on the corresponding median error obtained. The vertical dotted lines represent the expected error at each confidence level. Color code refers to the four CP models (*h(x)−g(x)*) employed. Markers refer to the requested confidence level.

Although the four CP models generally tend to have higher errors than expected, their median validity is close to the expected error for all models. Moreover, the CP models that use the XGB algorithm as their error model tend to have higher uncertainty than those that use the RF algorithm as their error model. For an 80% confidence level, the four CP models achieve the best uncertainty results, with values ranging from 1.6 to 2.1 pGI_50_ units. However, higher confidence levels are associated with higher uncertainty, so the choice of confidence level should be based on the specific task. pGI_50_ predictions at a lower confidence level are more reliable in this case.

Based on Figure 5, we explored the performance of CP models at an 80% confidence level since this level balances reliability and interpretability in the pGI_50_ predictions. CP models are compared to the non-CP models (i.e., a standard machine learning model). This comparison shows that, across all evaluated metrics (RMSE, Rp, and Rs), CP models consistently outperformed the non-CP models (Figure 4). More specifically, the median RMSE in the CP-valid predictions is about 0.3 pGI_50_ units lower than that of the non-CP predictions. This trend is also observed in correlation (Rp and Rs), with CP-valid predictions displaying a better alignment between the predicted and observed pGI_50_ values. One of the reasons for the superior performance of the CP models is their ability to assign uncertainty to their predictions. This is accomplished by generating prediction intervals associated with specific confidence levels. By doing so, the CP models inherently consider the uncertainties arising from distribution shifts, thus providing more reliable and accurate predictions.

Another significant result derived from Figure 6 is related to the CP-invalid predictions. Since CP-valid predictions exhibited superior results across all evaluated metrics (RMSE, RP, and Rs), compared to the broader non-CP predictions (non-CP = CP-valid U CP-invalid), it suggests that the errors in the CP-invalid predictions disproportionately contribute to the overall model inaccuracies. The concentration of errors within the CP-invalid predictions indicates potential challenges in accurately modeling uncertainties where the observed pGI_50_ values fall outside the predicted intervals.

**Figure 6.**
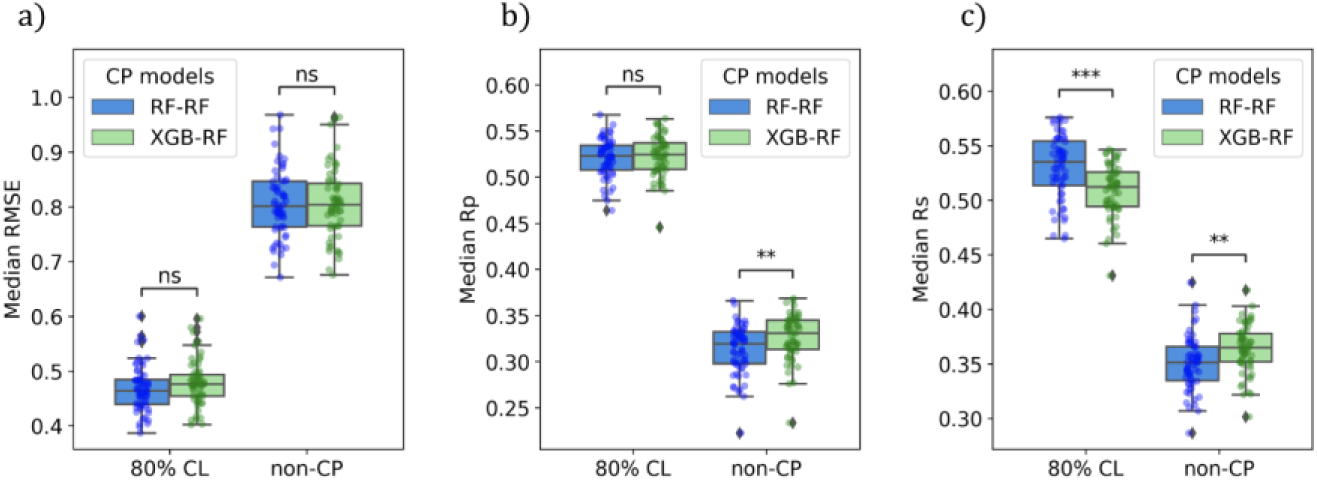
CP-valid predictions exhibit lower errors and higher correlations than non-CP predictions. RMSE, Rp, and Rs values were computed between the observed and predicted pGI_50_ values using either RF-RF or XGB-RF CP models. Each boxplot contains 60 points that represent the 7-fold-median a) RMSE, b) Rp, and c) Rs, summarizing their distributions across cell lines. Color code refers to the CP model used. The results of the statistical test performed for the models have been placed at the top of the boxplots. p-value annotation legend: ns: 5.00e-02 < p <= 1.00e+00; **: 1.00e-03 < p <= 1.00e-02; ***: 1.00e-04 < p <= 1.00e-03

Although there are no statistically significant differences in RMSE and Rp between the two CP models, a statistically significant difference exists in Rs. These differences are further explored to identify the strengths of each CP model. The RF-RF and XGB-RF models show comparable results in RMSE and Rp, indicating proficiency in reducing absolute errors and establishing linear relationships between predicted and observed pGI_50_ values. In addition, the RF-RF performs better in capturing non-linear dependencies, as shown by their Rs. These findings suggest that while the CP models may have similar performance in terms of overall prediction accuracy (RMSE) and linear correlations (Rp), they may differ in their ability to accurately capture the underlying patterns and non-linear relationships (Rs) among the 60 cell lines.

To highlight the proposed LDMO-CCP approach’s predictive accuracy, we compare its results against those reported in a previous study (34), in which we evaluated the performance of graph neural networks (GNNs), including the Directed Message Passing Neural Network (D-MPNN), on the same NCI-60 dataset. In Vishwakarma et al. (34), we benchmarked the standard ML versions of the RF and XGB models (i.e., non-CP models) and used a dissimilar-molecule split that aligns conceptually with the current clustering-based data partitioning approach.

While the best-performing D-MPNN model achieved slightly better RMSE values compared to the standard RF and XGB models, we demonstrate that our LDMO-CCP models not only offer well-calibrated uncertainty estimates but also yield lower RMSE scores than their best D-MPNN model (RMSE=0.585). This comparison underscores the practical value of CP-enhanced models and suggests that CP, when combined with a robust validation strategy like LDMO-CV and tree-based models, can match or exceed DL performance under realistic data splits.

Among the 60 cells, we identified both the best-performing and worst-performing cells based on their hit rates computed only over the CP-valid molecules. In cases where multiple cells had identical hit rates, we utilized uncertainty as a tie-breaking measure. Table 1 shows these results. When considering only CP-valid molecules, both the RF-RF model and the XGB-RF model achieve a 100% hit rate, which highlights the importance of using CP to calibrate the predictions. The RF-RF model identifies cell line SK-OV-3 as the optimal choice, while the XGB-RF model selects cell line TK-10. Based on the uncertainty, the RF-RF assigns more confidence in the predictions for the molecules tested against the SK-OV-3 cell line.

**Table 1.**
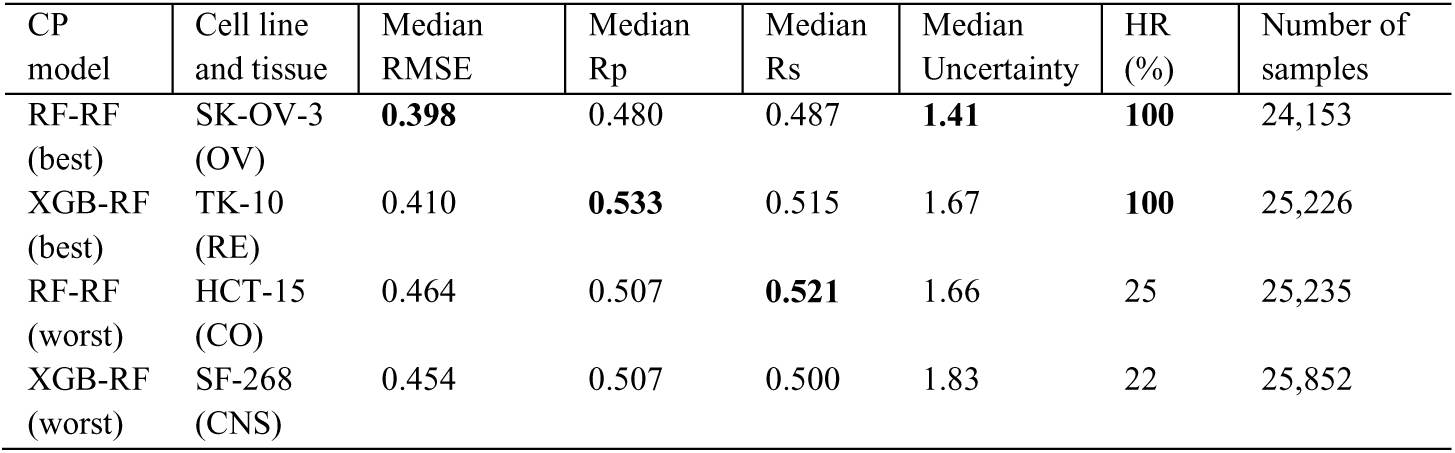
Performance summary of best– and worst-performing cell line (CP models) in terms of the out-of-sample hit rate. The predictions come from either the RF-RF or XGB-RF modes, using MFPs+PCDs as the input feature set. The models were trained and evaluated using LDMO CV, with the reported values representing the median scores obtained across the folds. The evaluation metrics include median RMSE/Rp/Rs, median uncertainty, and hit rate (HR). These metrics are computed only for the CP-valid molecules at an 80% confidence level. The hit rate is obtained by applying a threshold of 6 pGI_50_ units that distinguishes between active and inactive molecules. The last column indicates the total number of CP-valid molecules in the dataset used for the calculation of the performance metrics. In bold are highlighted the best-performing values for each metric.

| CP model | Cell line and tissue | Median RMSE | Median Rp | Median Rs | Median Uncertainty | HR (%) | Number of samples |
| --- | --- | --- | --- | --- | --- | --- | --- |
| RF-RF (best) | SK-OV-3 (OV) | **0.398** | 0.480 | 0.487 | **1.41** | **100** | 24,153 |
| XGB-RF (best) | TK-10 (RE) | 0.410 | **0.533** | 0.515 | 1.67 | **100** | 25,226 |
| RF-RF (worst) | HCT-15 (CO) | 0.464 | 0.507 | **0.521** | 1.66 | 25 | 25,235 |
| XGB-RF (worst) | SF-268 (CNS) | 0.454 | 0.507 | 0.500 | 1.83 | 22 | 25,852 |

### 4.3 Useful predictions are achieved despite training on highly dissimilar molecules

To simulate a real screening scenario, we focused on the best-performing cell lines predicted by the two CP models, SK-OV-3 and TK-10 (see Table 1 for performance metrics). We first identify the 100 CP-valid molecules with the highest predicted pGI50 values for these cell lines. Subsequently, we narrowed our search to the three molecules with the highest measured pGI50 values from this subset. This mimics the real screening scenario, where a large pool of candidate molecules is initially considered, followed by a refinement process to identify the most promising candidates for further analysis. Our goal is to measure how dissimilar the best CP-valid molecules are from the molecules the CP model already saw during the training stage. To accomplish this, we search for its closest counterpart for each of the three best-performing CP-valid molecules by computing the Tanimoto similarity within the training clusters. This search yields six candidates (one per cluster, as the test cluster is not considered during this search), from where we identify the training molecule that exhibits the highest degree of similarity to the best-selected CP-valid molecule. This selection is made as the other five candidates exhibit a higher dissimilarity to this particular molecule.

Table 2 shows that regardless of the CP model used, either RF-RF or XGB-RF, all the similarities observed between the molecules were relatively small, ranging from 0.29 to 0.42. These modest similarity values indicate subtle structural similarities or shared features among the molecules (Figure 7). Using the XGB-RF model, the three most promising molecules identified were predicted with pGI_50_ values greater than 6; therefore, these molecules are considered active. Notably, the most similar molecule found in the training set for all three promising molecules was the same (NSC-662190), highlighting a consistent pattern in their structural similarities. In contrast, among the three promising molecules obtained using the RF-RF model, only one was considered an active molecule (predicted pGI_50_=6.8).

**Figure 7.**
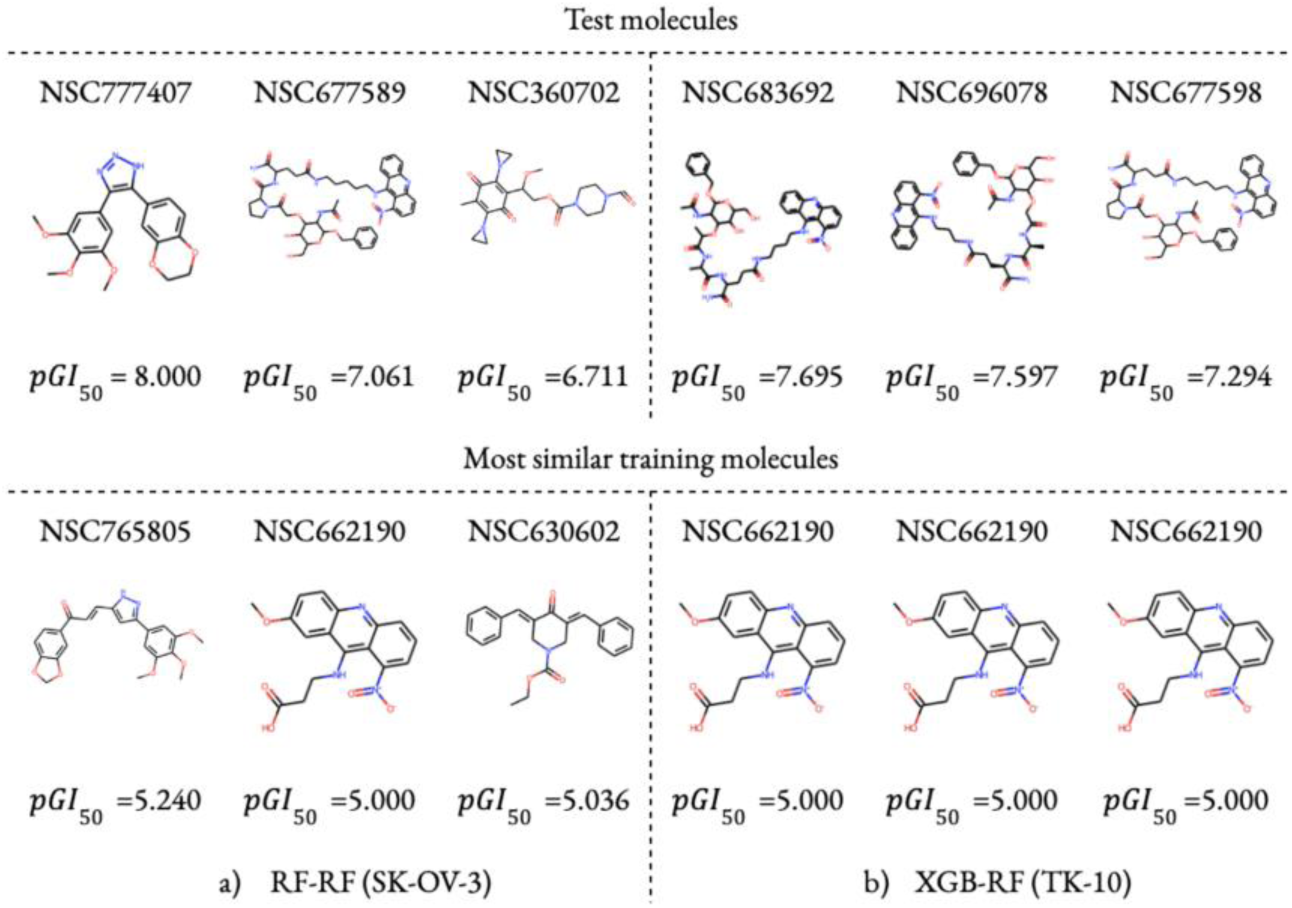
Chemical structures of the three molecules with the highest measured pGI50s among the top 100 CP-valid predictions in the test set along with their most similar training molecules (non-outlier dataset). Test molecules are shown at the top, while their most similar training molecules are shown at the bottom. The measured pGI_50_ is shown at the bottom of each molecule. Regardless of the CP model used, either a) RF-RF or b) XGB-RF, all the similarities observed between the molecules were relatively small, ranging from 0.29 to 0.42. See Table 2 for metrics performance.

**Table 2.** Similarity analysis between the best-performing CP-valid molecules and the training molecules (non-outlier dataset). For the best-performing cell lines, SK-OV-3 and TK-10 (see Table 1 for performance metrics), we mimic the real screening scenario, where a large pool of candidate molecules is initially considered, followed by a refinement process to identify the most promising candidates for further analysis. The three molecules with the highest measured pGI_50_s among the top 100 CP-valid predictions in the test set are shown in this table (thus simulating the results of *in vitro* testing these 100 molecules). For each of these three CP-valid molecules, we search for their respective 6 closest counterparts within the training clusters, from where we identify the training molecule that exhibits the highest degree of similarity to the best-selected CP-valid molecule. The cell line, NSC-ID, measured pGI_50_, and predicted pGI_50_ are included in this table

| CP model | Cell line | Test molecule |  |  | Most similar training molecule |  |  |
| --- | --- | --- | --- | --- | --- | --- | --- |
|  |  | NSC ID | Measured pGI<sub>50</sub> | Predicted pGI<sub>50</sub> | NSC ID | Measured pGI<sub>50</sub> | Similarity |
| RF-RF | SK-OV-3 | 777407 | 8.000 | 5.481 | 765805 | 5.240 | 0.422 |
| RF-RF | SK-OV-3 | 677598 | 7.061 | 6.815 | 662190 | 5.000 | 0.317 |
| RF-RF | SK-OV-3 | 360702 | 6.711 | 5.326 | 630602 | 5.036 | **0.290** |
| XGB-RF | TK-10 | 683692 | 7.695 | 7.026 | 662190 | 5.000 | 0.345 |
| XGB-RF | TK-10 | 696078 | 7.597 | 6.904 | 662190 | 5.000 | 0.339 |
| XGB-RF | TK-10 | 677598 | 7.294 | 7.109 | 662190 | 5.000 | 0.317 |

### 4.4 CP predictions with low uncertainty are more accurate than those with high uncertainty

The LDMO-CCP models detailed in the previous sections underwent training and evaluation within a relatively controlled environment represented by the non-outlier dataset. However, a question remains concerning the ability of a CP model to effectively generalize to molecules with dissimilar chemical structures from those in its training set. To address this, we retrain an LDMO-CCP model on the non-outlier datasets and test it on the outlier dataset for each of the 60 cell lines. Thus, this experimental design encompasses a comprehensive evaluation of the adaptability and performance of a CP model in handling different scenarios to which it was exposed during training. The two best CP models are re-trained per cell line using the complete set of non-outlier molecules, for which 6 of the 7 clusters are used as the proper training set, and the remaining cluster is used as the calibration set (Fig. 3-a).

The results from the XGB-RF and RF-RF models agree in identifying NCI-H322M as the cell with the highest predicted hit rate, demonstrating consensus between the two models. In the subsequent analysis, we compared the results of the 5% of molecules with the lowest uncertainty to the 5% with the highest uncertainty. It is important to note that by considering only these predicted uncertainties, this analysis does not employ any information whatsoever from the measured pGI_50_ values in the independent test sets (outlier molecules tested on each cell line). This is suitable for prospective studies, where we are interested in making predictions or drawing conclusions about future outcomes based on data that has not yet been generated.

Figure 8 shows that both the RF-RF and XGB-RF models benefit when considering only the 5% of molecules with the lowest uncertainty. In this scenario, the RF-RF model achieves lower RMSE while the XGB-RF achieves higher Rp and Rs. In contrast, for the 5% of molecules with the highest uncertainty, the RF-RF model shows better Rs but a higher RMSE compared to those obtained by the XGB-RF CP model. These findings underscore the significance of considering uncertainty levels in model predictions, as lower uncertainty is consistently associated with enhanced predictive accuracy and reliability across both models.

**Figure 8.**
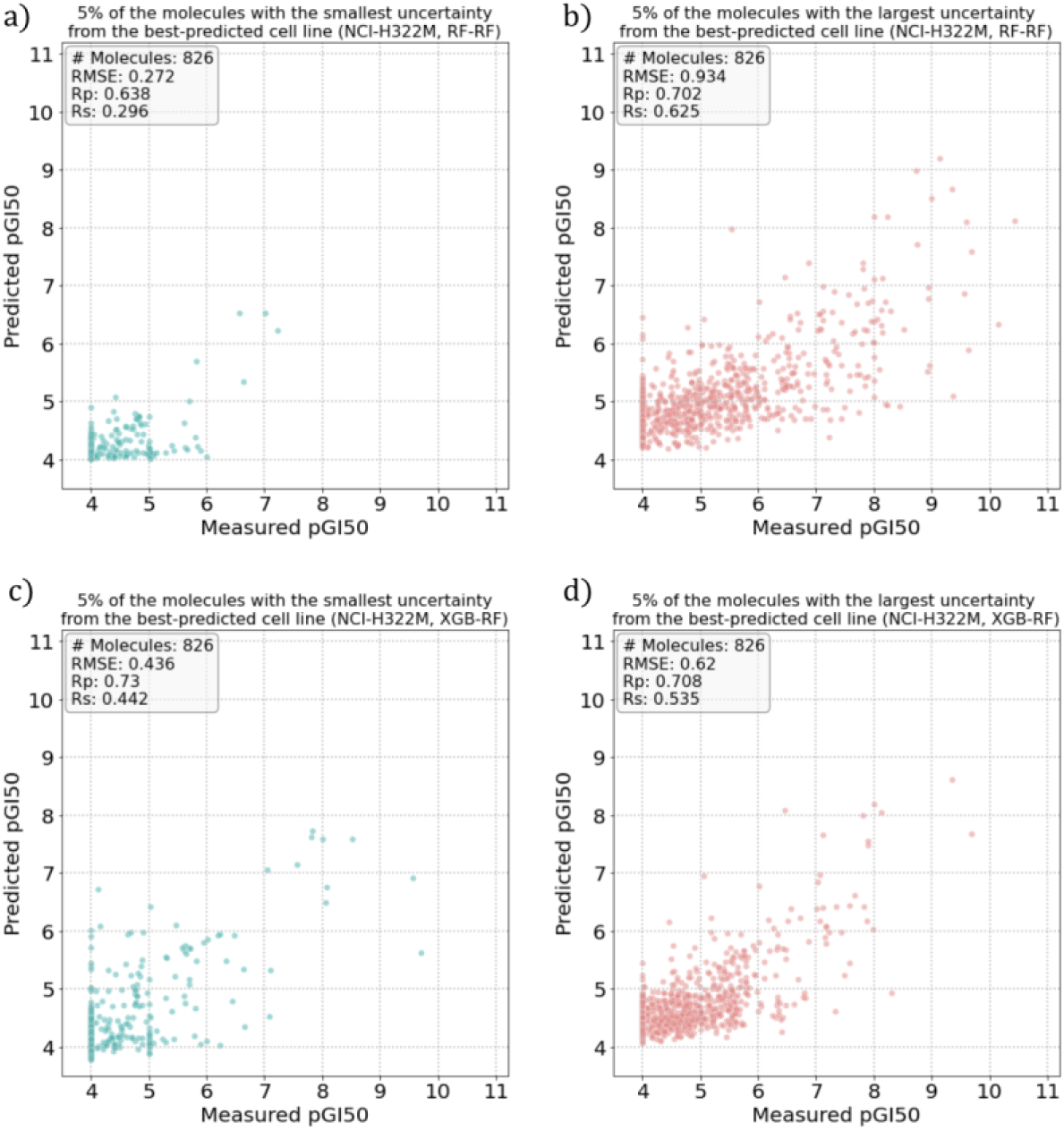
The prediction of the pGI_50_ of molecule-cell line pairs improves when CP is used. Observed and predicted pGI_50_ value in the best-predicted cell line, obtained by the RF-RF (top) and the XGB-RF (bottom) CP models. Predictions are obtained at an 80% confidence level. The uncertainty is calculated by multiplying the predicted error for a given molecule by the non-conformity score chosen at an 80% confidence level. Each plot contains the 5% (826 molecules) of the test set molecules, either those with the smallest (left) or largest uncertainty (right). The anomalous density of molecules at pGI_50_=4 is due to the inhibitory activity of weaker molecules being rounded up at the closest considered pGI_50_ of 4.

### 4.5 The ensemble of CP models can predict the NCI-60 cell line selectivity of a test molecule

An inherent advantage of the CP framework is its ability to assign a degree of uncertainty at an instance level. We leverage this characteristic to identify the best-performing molecules among the 60 cell lines based on their respective degrees of uncertainty for both CP models. In this instance-level evaluation, molecules with lower uncertainty are considered the best-performing. The uncertainty is calculated by multiplying the predicted error for a given molecule by the non-conformity score chosen at a confidence level. In essence, the uncertainty corresponds to the length of the predicted interval and reflects the range within which the observed value of the molecule is likely to fall.

To achieve this task, we first merge the data for each molecule across all 60 cell lines and define additional criteria to keep only the CP-valid and most potent molecules (pGI_50_ >= 6). After this filter, the RF-RF model retained 5,252 NSC-Cell points for 301 small molecules screened against the 60 cancer cell lines, resulting in a matrix completeness of 29.08%. We further narrowed our search to only those NSCs where data is available for at least 50 cell lines. As a result, we identified 37 NSC, from which we selected the best-performing CP-valid molecule based on their uncertainty. Similarly, for the XGB-RF model, the filtering process yielded 6,075 NSC-Cell points for 567 small molecules, representing a matrix completeness of 17.86%. Subsequently, we identified 20 NSCs for which the data is available for at least 50 cell lines, and from which we selected the best-performing CP-valid molecule based on their uncertainty.

Figure 9 shows that the molecule NSC-ID: 747457, identified by the RF-RF CP model, demonstrates a relatively confident prediction (uncertainty of 1.8 units) and high potency across 52 cell lines, with pGI_50_ values higher than 6. In contrast, the XGB-RF CP model identified the molecule NSC-7521 (proscillaridin), which demonstrates a relatively confident prediction given by an uncertainty of 1.3 units. Moreover, this molecule demonstrates a high potency across an even broader spectrum of 58 cell lines, with pGI_50_ values higher than 8. Therefore, in terms of uncertainty and potent activity, the XGB-RF CP model, with the proscillaridin molecule, outperforms the results from the RF-RF CP model.

**Figure 9.**
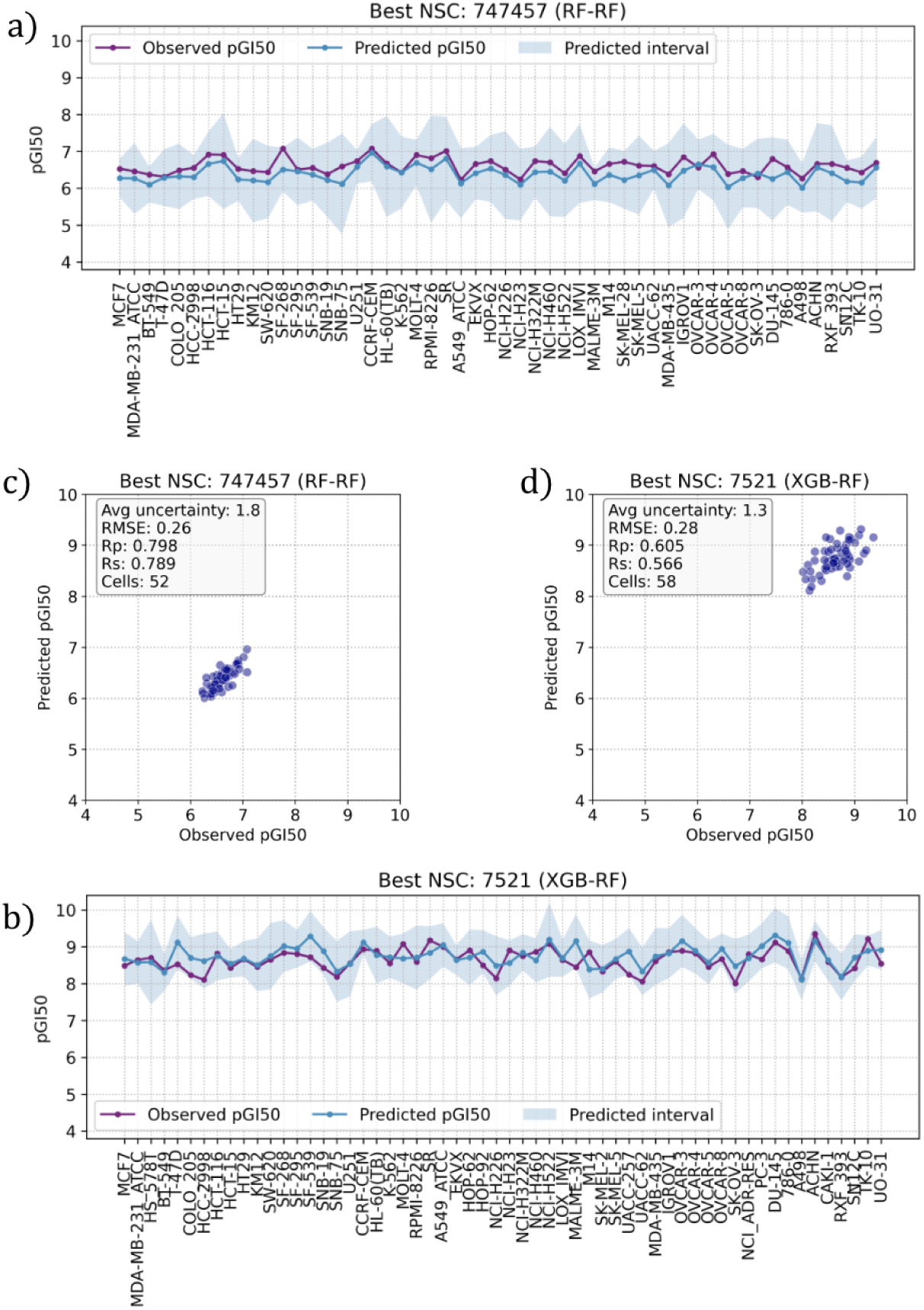
The ensemble of CP models can predict the selectivity of a dissimilar molecule across NCI-60 cell lines. Relationship between the measured and predicted pGI_50_ values across different cells, for the best-performing 80% CP-valid predictions. The analysis prioritizes the identification of molecules predicted reliable (CP-valid) and potent pGI_50_ ≥ 6, either by the (top) RF-RF or (bottom) XGB-RF model. The best CP-valid molecule (NSC:747457) is selected from a pool of 37 molecules for the RF-RF, while for the XGB-RF, the best CP-valid molecule (Proscillaridin; NSC:7521) is selected from a pool of 20 molecules tested on at least 50 cell lines. The best-performing molecule is based on its uncertainty (length of the interval predicted). The uncertainty is calculated by multiplying the predicted error for a given molecule by the non-conformity score chosen at an 80% confidence level. The evaluation metrics also include RMSE, Rp, and Rs. Each dot in the plot corresponds to a cell line treated with the corresponding molecule. To facilitate comparison, the y-axis in all figures is set to the same scale.

## 5 Discussion

This study introduces the LDMO-CCP approach, a modified version of CCP that utilizes molecular clustering as the data partitioning strategy. This modification aims to improve the model’s generalization capacity across various chemical spaces, especially when working with structurally diverse compounds. The proposed LDMO-CCP models consistently outperformed standard ML models in predicting the inhibitory activity of molecules on each NCI-60 cancer cell line. In contrast to traditional ML models, CP offers the additional advantage of generating prediction sets with a calibrated level of uncertainty, thereby ensuring that confidence metrics are associated with predictions. This feature is particularly beneficial in drug design, where decisions are made under uncertainty, and incorrect predictions can impact further stages.

Compared to DL methods such as the D-MPNN (34), which leverages dropout strategies to enhance generalization, the proposed LDMO-CCP approach demonstrates superior performance while requiring significantly less intensive hyperparameter tuning. Notably, CP-valid predictions derived from the LDMO-CCP models outperform D-MPNN in terms of predictive accuracy under dissimilar data splits while also providing reliable uncertainty estimates.

Identifying Proscillaridin (NSC: 7521) using the proposed LDMO-CCP approach underscores the promise of CP-based models in early-phase drug design. Proscillaridin emerges as a potential inhibitory compound, showing broad-spectrum activity across multiple cancer cell lines in the NCI-60 panel. Moreover, proscillaridin has been found to effectively induce oxidative stress and apoptosis in the cell line A549 (45), showcasing its potential as an anti-cancer agent and, in turn, providing support for the proposed LDMO-CCP approach. Traditional ML models may not have captured this result with the same reliability at the instance level, as they lack the uncertainty quantification layer that CP models provide. This quantification is essential due to the time and cost involved in experimental validation. CP-based models can effectively minimize the risk of both false positives and false negatives.

Additionally, while our results are promising, further prospective validation is required to confirm the utility of these models in real-world settings. Leveraging advanced graph-based models for molecular representation can be explored to improve the accuracy when predicting the inhibitory activity of molecules (46,47). By merging the generalization ability of DL (48,49) with the formal reliability assurances of CP, we could improve performance on out-of-distribution data. Moreover, recent research has effectively utilized CP frameworks in ultra-large chemical libraries (50), showcasing their scalability and practicality in real-world virtual screening situations with millions of compounds. These results confirm the appropriateness of CP for extensive drug discovery initiatives.

## 6 Conclusions

We have developed CP models for out-of-sample prediction and confidence intervals of the pGI50s of molecules on each NCI-60 cell line. This panel contains data from 60 distinct human tumor cell lines and is a comprehensive resource for identifying and characterizing novel compounds with growth inhibition or killing of tumor cell lines on various cancer types.

The LDMO-CV approach used to build and evaluate these CP models on clusters of non-outlier molecules leads to models that generally perform well on unseen molecules dissimilar to those in the training. This is the most common scenario as real-world compound libraries contain many more molecules with higher diversity than those already tested on the cell line. We observed that the higher the requested confidence level, the higher the prediction uncertainty becomes, which in turn results in worse RMSE when predicting such dissimilar molecules. CP models trained on all non-outlier molecules and tested on outlier molecules further establish the ability of these models to handle training-test partitions introducing strong distribution shifts.

Overall, we showed that predictions with associated confidence intervals enhance decision-making in that novel molecules with smaller prediction errors can be selected for further exploration and experimental validation. Furthermore, these CP models could predict the selectivity of dissimilar molecules across NCI-60 cell lines. Focusing on the 80% CP-valid predictions, we identified molecules exhibiting both reliability and potency. Notably, NSC:747457 and NSC:7521 (Proscillaridin) emerged as the CP-valid molecules whose cell line selectivity could be predicted best, highlighting the potential of this approach for strategic molecule prioritization in drug discovery efforts.

## Author Contributions

P.J.B. conceived and led the study. S.H-H. wrote the code to run the experiments. All authors analyzed the results from the experiments. P.J.B. wrote the paper with assistance from S.H-H. and Q.G.

## Acknowledgment

This work was supported by CONAHCYT [S.H-H.], and the Wolfson Foundation and the Royal Society for a Royal Society Wolfson Fellowship [P.J.B.].

## Data availability

The data used in our study are publicly available in NCI-60 (https://wiki.nci.nih.gov/display/NCIDTPdata/NCI-60+Growth+Inhibition+Data), whereas all the partitions made for this study can be found at GitHub (https://github.com/Sahet11/nci60_clustering).

## Code availability

The source code for replicating this study’s results is freely available at GitHub (https://github.com/Sahet11/STL_LDMO_project).

